# Determining the pathogenicity of variants of uncertain significance and identification of a founder variant in the epilepsy-associated gene, *SZT2*

**DOI:** 10.1101/2021.01.06.425612

**Authors:** Jeffrey D Calhoun, Miriam C Aziz, Hannah C Happ, Jonathan Gunti, Colleen Gleason, Najma Mohamed, Kristy Zeng, Meredith Hiller, Emily Bryant, Divakar Mithal, Irena Bellinski, Lisa Kinsley, Mona Grimmel, Eva MC Schwaibold, Constance Smith-Hicks, Anna Chassevent, Marcello Scala, Andrea Accogli, Annalaura Torella, Pasquale Striano, Valeria Capra, Lynne M. Bird, Issam Ben-Sahra, Nina Ekhilevich, Tova Hershkovitz, Karin Weiss, John Millichap, Elizabeth E Gerard, Gemma L Carvill

## Abstract

Biallelic pathogenic variants in *SZT2* result in a neurodevelopmental disorder with shared features, including early-onset epilepsy, developmental delay, macrocephaly, and corpus callosum abnormalities. SZT2 is as a critical scaffolding protein in the amino acid sensing arm of the mTOR signaling pathway. Due to its large size (3432 amino acids), lack of crystal structure, and absence of functional domains, it is difficult to determine the pathogenicity of *SZT2* missense and in-frame deletions. We report a cohort of twelve individuals with biallelic *SZT2* variants and phenotypes consistent with *SZT2*-related neurodevelopmental disorder. The majority of this cohort contained one or more *SZT2* variants of uncertain significance (VUS). We developed a novel individualized platform to functionally characterize *SZT2* VUSs. We identified a recurrent in-frame deletion (SZT2 p.Val1984del) which was determined to be a loss-of-function variant and therefore likely pathogenic. Haplotype analysis determined this single in-frame deletion is a founder variant in those of Ashkenazi Jewish ancestry. Overall, we present a FACS-based rapid assay to distinguish pathogenic variants from VUSs in *SZT2*, using an approach that is widely applicable to other mTORopathies including the most common causes of the focal genetic epilepsies, *DEPDC5, TSC1/2, MTOR* and *NPRL2*/3.

## Introduction

*SZT2 (<u>S</u>ei<u>z</u>ure <u>T</u>hreshold 2)* encodes a large (>350 kDa) protein with ubiquitous tissue expression and no homology to known functional domains ^1^. The first association of *SZT2* with seizure susceptibility emerged from a mutagenesis screen in mice ^1^. In this study, homozygous *Szt2* knockout mutant mice seized at a lower electrical input relative to wildtype littermates in an acute electroconvulsive model ^1^. Years later, biallelic *SZT2* pathogenic variants were identified in a small cohort of patients with infantile-onset epilepsy and dysgenesis of the corpus callosum ^2^. Since then, several small case studies report biallelic *SZT2* variants in association with a neurodevelopmental disorder (NDD) characterized primarily by early-onset focal epilepsy, developmental delays, macrocephaly and corpus callosum abnormalities ^2-16^.

For years the function of *SZT2* remained elusive until 2017 when *SZT2* was identified as a part of the KICSTOR complex, a required component of the amino acid sensing arm of the mTORC1 pathway ^17; 18^. The KICSTOR complex, including SZT2, localizes to the lysosomes only in the presence of amino acids in the extracellular environment ^17^. In genome-edited cells lacking endogenous expression of *SZT2*, mTORC1 activity no longer depended on amino acids, i.e. mTORC1 was constitutively active ^17^. Moreover, cells lacking *SZT2* exhibited amino acid-insensitive localization of mTOR to the lysosomal surface, suggesting that SZT2 is a key scaffolding protein ^17^. Based on these two pivotal studies, we hypothesize that biallelic *SZT2* loss-of-function (LoF) variants produce a mTORopathy due to constitutive mTORC1 activity ^17; 18^.

The majority of the pathogenic *SZT2* variants described to date are truncations, resulting in complete *SZT2* LoF (null). However, there are now also many cases of both missense variants and in-frame deletions that are more difficult to classify, driven primarily by our inability to determine the effect of single amino acid alterations on SZT2 function. We focus primarily on these variants of uncertain significance (VUS) and devise a functional assay to identify LoF SZT2 alleles. Moreover, we use this approach to demonstrate that a recurrent single amino acid deletion is both likely pathogenic and a founder variant in those of Jewish ancestry.

## Subjects and Methods

### Study participants

Individuals with candidate causative *SZT2* variants were identified by clinical exome sequencing (n=11) or gene panel (n=1). The individuals at Northwestern Memorial Hospital and Lurie Children’s Hospital were consented to research under an IRB approved study. We obtained de-identified genetic and clinical data from external colleagues for cases identified through Genematcher ^19^. These individuals were consented for research under IRB approved studies at their local institutions. *SZT2* variant classification was performed according to the American College of Medical Genetics (ACMG) criteria ^20^.

### Haplotype analysis in individuals with the recurrent p.Val1984del variant

Exome sequencing data from probands and parents with the *SZT2* p.Val1984del variant were obtained. Variant call files (vcfs) were generated with standard GATK best practices. Briefly, sequencing reads were aligned to the human genome (hg38) with BWA-MEM followed by calling and genotyping alleles with GATK HaplotypeCaller and GATK GenotypeGVCFs. Then, GATK SelectVariants was used to subset a vcf file containing variants within 20 Mb of *SZT2* p.Val1984del in the individual homozygous due to uniparental disomy (UPD) of all of chromosome 1 (proband 3). SNVs and indels in this region were filtered by minor allele frequencies in gnomAD (see Web Resources) to generate a list of candidate variants for haplotype analysis. Segregation of alleles in independent trios was used to define haplotype boundaries.

### Generation of gene-edited cell lines

pSpCas9(BB)-2A-Puro (PX459) V2.0 was a gift from Feng Zhang (Addgene plasmid # 62988; http://n2t.net/addgene:62988; RRID:Addgene_62988). gRNAs targeting SZT2 exons (Table S1) were cloned into PX459 as previously described ^21^. HEK 293T (ATCC® CRL-3216™) cells were seeded into 24-well plates and transfected with pX459 plasmid with cloned gRNAs (500-1000 ng) and ssODN repair oligo (1-2 uL of 10 μM stock) with lipofectamine 3000 (Invitrogen# L3000001) according to manufacturer’s instructions. Cells were treated with puromycin (2.5 μg/mL) for 48 hours beginning the day after transfection. Cells were replica plated for cryopreservation and genomic DNA isolation with PureLink™ Genomic DNA Mini Kit (Invitrogen K182002) according to manufacturer’s instructions. gRNA targeting of SZT2 exon was confirmed by T7 endonuclease I (NEB# M0302) according to manufacturer’s instructions. Homology-directed repair was confirmed by amplicon sequencing. Individual clones were collected by limited dilution cloning in 96 well plates followed by similar replica plating as described above. Genomic DNA from each clone was screened by Sanger sequencing as well as amplicon sequencing to confirm genotype as either (1) homozygous for the homology-directed repair (HDR) allele or (2) compound heterozygous for the HDR allele and an out-of-frame indel predicted to lead to *SZT2* LoF. All primer sequences provided in Table S1.

### Amplicon sequencing

Amplicons with Illumina adaptors were generated by two rounds of PCR including the introduction of unique barcodes and standard Illumina primers. Amplicons were sequenced to a depth of at least 1,000X on an Illumina Miniseq according to manufacturer’s recommendation. Alleles were analyzed with CRISPResso2 using the web interface or command line workflow ^22^.

### Immunoblot analysis of mTORC1 activity

Amino acid starvation was performed as previously described ^17^. Briefly, cells plated in poly-L-lysine coated plates were rinsed in Dulbecco’s phosphate-buffered saline (DPBS) (Gibco# 14190250) twice before the addition of amino acid-free Dulbecco’s minimal essential media (DMEM) containing 10% dialyzed FBS. For the amino acid starved condition, cells were starved of amino acids for 60 min at 37 °C. For cells treated with amino acids, cells were starved of amino acids for 50 min at 37 °C followed by incubation with amino acid containing DMEM for 10 min at 37 °C. Cells were briefly rinsed with ice-cold DPBS twice. Cells were scraped in ice-cold DPBS and pelleted at 300 xg for 5 min. The cells were lysed in RIPA Lysis and Extraction Buffer (Thermo Scientific 89900) supplemented with EDTA-free protease inhibitors (Roche Complete PI EDTA-free; Sigma 11836170001) and PhosSTOP™ phosphatase inhibitors (Sigma 4906845001) for 30 min at 4 °C. Insoluble material was removed by pelleting at 12,000 xg for 20 min. Protein concentration was determined by BCA assay (Pierce™ BCA Protein Assay Kit; Cat# 23225) and equal amounts (10-50 μg) of protein were loaded for SDS-PAGE (4-12% gradient gel). Proteins were transferred to PVDF membrane and blocked for 60 min at room temperature (RT) with 5% bovine serum albumin (BSA) in PBS-T (PBS + 0.05% Tween20). Primary antibodies (Table S2) were incubated overnight at 4 °C in blocking buffer. Membranes were washed 3x 5 min in PBS-T prior to incubation with secondary antibodies (Table S2) for 60 min at RT. After washing 4x 5 min in PBS-T, membranes were incubated in Amersham ECL Prime (GE Healthcare) according to manufacturer’s instructions. Membranes were imaged on a Licor Odyssey Fc for 30 sec to 60 min.

### FACS analysis of mTORC1 activity

Amino acid starvation was performed as described above. After starvation, cells were washed twice with DPBS supplemented with 1% v/v Phosphatase Inhibitor Cocktail 3 (Sigma; P0044). Cell were trypsinized using trypLE (Gibco) supplemented with 1% v/v Phosphatase Inhibitor Cocktail 3. Cells were pelleted (300 x g for 5 min at RT) and resuspended in BD Fixation/Permeabilization solution (BD Biosciences 554714). After incubation on ice for 20 min, cells were pelleted and washed twice with BD Permeabilization/Wash buffer. Phosphorylated S6 (p-S6) was labeled with Alexa488 conjugated antibody (Table S2) for 30 min on ice followed by two washes with BD Permeabilization/Wash buffer. Cells were flow sorted on the BD FACSMelody 3-laser cell sorter. At least 200,000 cells were collected for cells with dim Alexa488-signal (P-S6^LOW^) and those with bright Alexa488-signal (P-S6^HIGH^). Genomic DNA was prepared by manufacturer’s recommendation using PureLink™ Genomic DNA Mini Kit (Invitrogen K182002) with slight modification to include a 40 min incubation at 90 °C to reverse crosslinks. Amplicon sequencing and Sanger sequencing were performed on unsorted and sorted cells to determine the <u>c</u>onstitutive <u>m</u>TORC1 <u>a</u>ctivity <u>s</u>core (CMAS). CMAS is calculated for each allele present in the amplicon sequencing dataset. CMAS = % alleles in P-S6^HIGH^ / % alleles in unsorted cells. CMAS represents an enrichment score to determine whether cells with high mTORC1 activity are enriched for proband-derived SZT2 missense alleles (HDR-mediated). Allele percentage derived from amplicon sequencing and used to calculate CMAS are reported in Table S3.

## Results

### Genetic characterization of individuals with bi-allelic SZT2 variants

We describe 12 individuals with bi-allelic *SZT2* variants and of these 24 alleles, truncating variants were identified in a quarter (6/24) of the cohort (Figure S1; Table 1). This included one individual (Individual 2) who had bi-allelic *SZT2* truncating variants^4^, and five individuals carrying a truncating variant on one allele and a single amino acid change on the other (Individuals 1, 5, 6, 7, 8). Most individuals carried at least one *SZT2* VUS, and these variants accounted for the majority of the variants present in the cohort (missense: 12/24, 50%; in-frame del: 5/24, 21%). One variant (p.Val1984del) was identified in multiple individuals in our cohort and has been previously reported in a single individual ^11^.

**Table 1.**
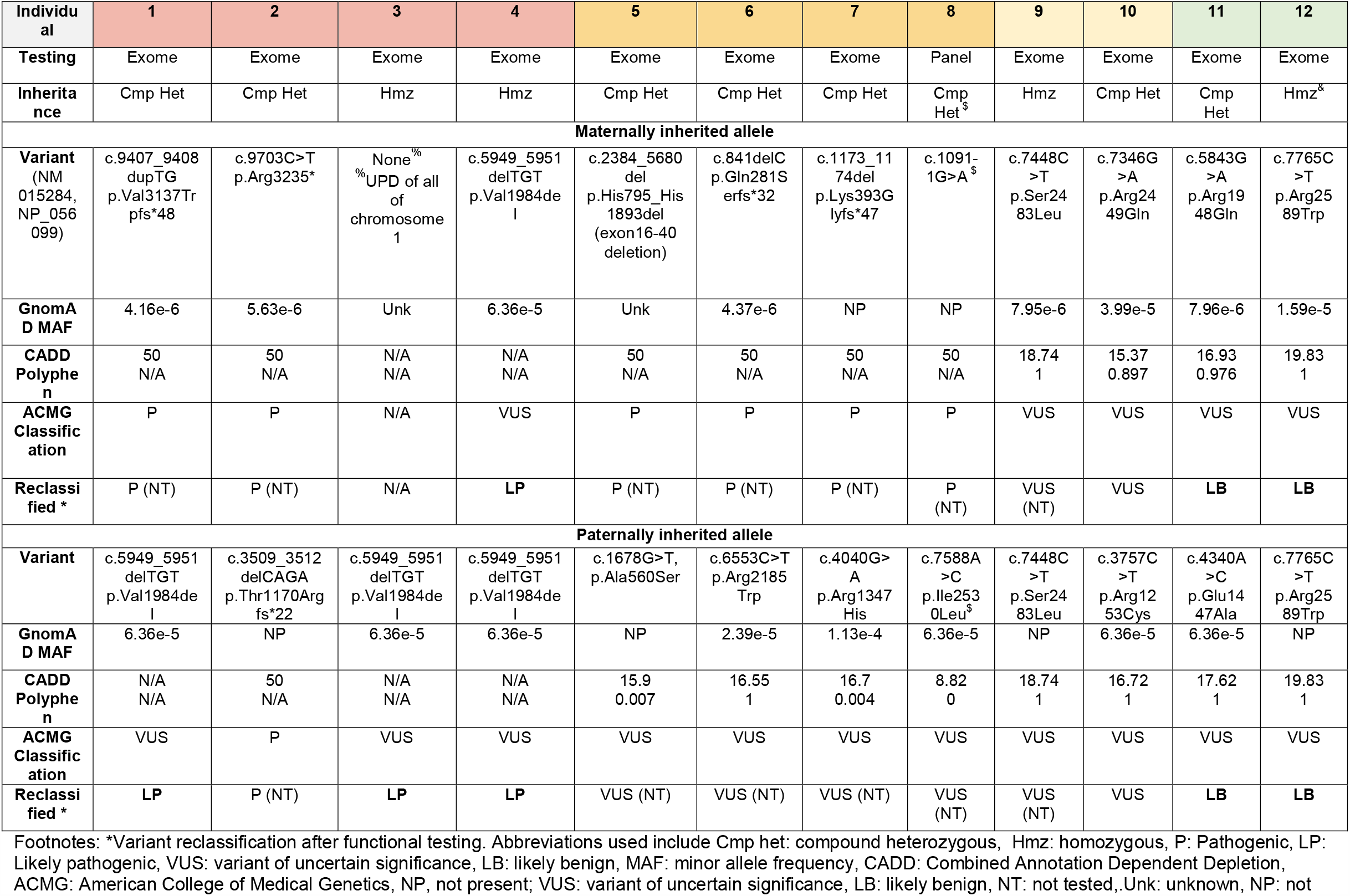
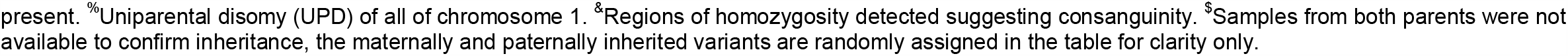
Genetic details of biallelic *SZT2* variants.

### Effect of patient-specific SZT2 variants on mTORC1 activity

We designed a functional assay to measure the effect of these *SZT2* VUSs. We utilized CRISPR/Cas9 to edit HEK293T cells at the endogenous *SZT2* locus and then quantified mTORC1 signaling in cells starved or starved and subsequently treated with amino acids (Figure 1A). Notably HEK293T are diploid for chromosome 1 on which SZT2 is located. First, we created a HEK293T SZT2 null cell line (*SZT2*^KO/KO^) and recapitulated the constitutive mTORC1 activity and insensitivity to the presence of amino acids previously observed in *SZT2* null cell lines ^17; 18^ (Figure S2).

**Figure 1.**
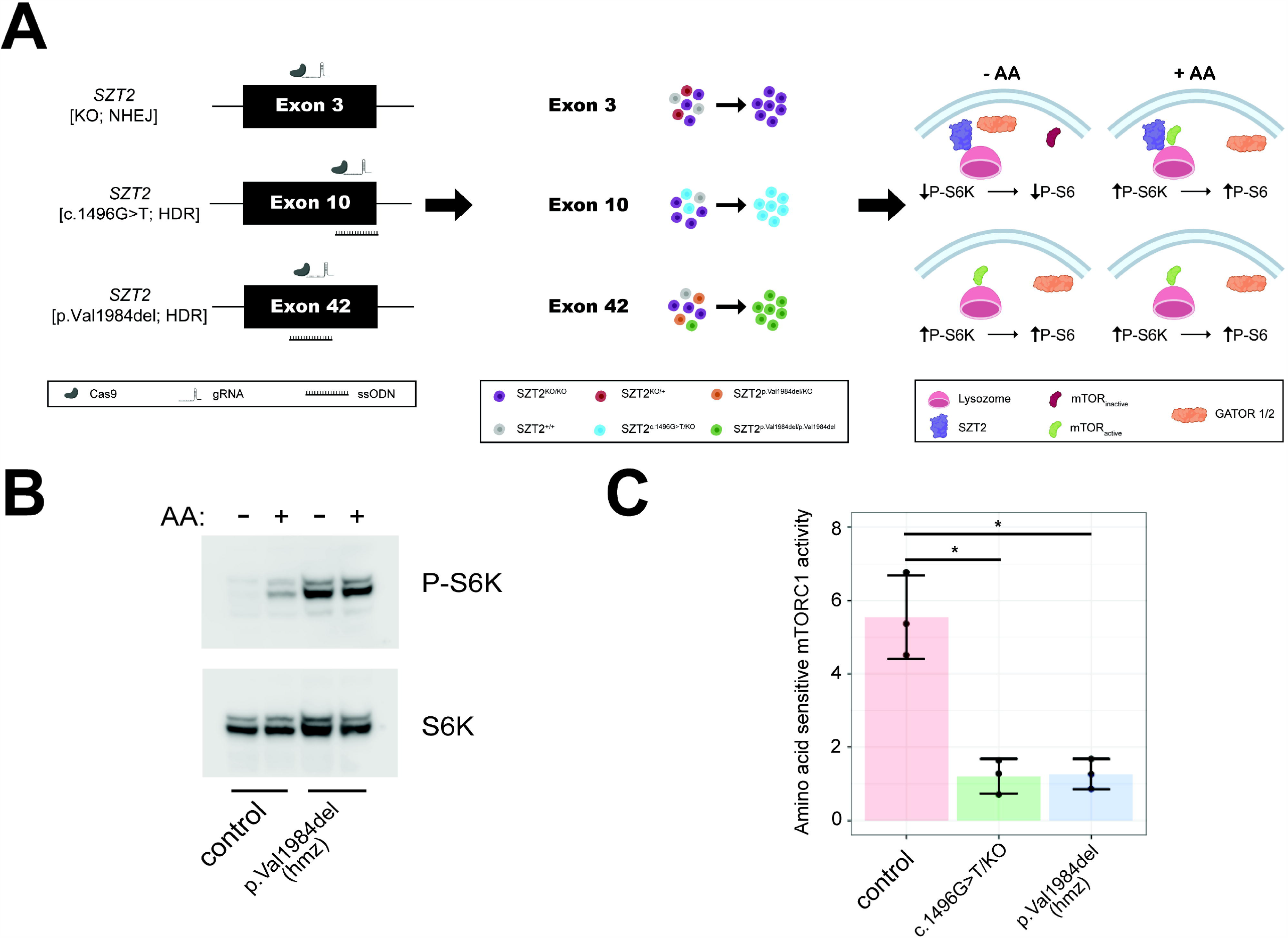
Development of gene editing approach for functional characterization of *SZT2* VUSs. (A) Individual cell clones either homozygous or heterozygous (+ LoF on other allele) for an individual *SZT2* variant were generated by gene editing followed by limiting dilution cloning. Immunoblot for amino acid sensitive mTORC1 activity by P-S6K levels was used to determine whether individual variants caused *SZT2* loss of function. Homology-directed repair rate of cells generated by transfection of px459 encoding targeting gRNAs was analyzed using amplicon sequencing and varied between 12-25%. (B) Immunoblot of mTORC1 activity in control HEK cells and homozygous HEK SZT2^p.Val1984del/p.Val1984del^ clone. (C) Densitometric quantification of (B). At least 3 individual replicates were performed. * = p < 0.05 (T-test).

We examined one previously published likely pathogenic *SZT2* variant (c.1496G>T) and the recurrent p.Val1984del VUS identified in three patients in our cohort (Probands 1,3,4) along with one individual described previously ^2; 11^. c.1496G is the last nucleotide of exon 10 and c.1496G>T yields two possible transcripts: (1) exon skipping resulting in *SZT2* p.Gly412Alafs*86 or (2) missense variant *SZT2* p.Ser499Ile. RT-PCR from patient fibroblasts suggested exon 10 skipping, though it was unclear whether any of the predicted missense allele p.Ser499Ile transcript was generated, and if so, whether this missense VUS impacted *SZT2* function^2^. We generated a compound heterozygous *SZT2* cell line (HEK.*SZT2*^c.1496G>T/KO^) and by analysis of RNA transcripts determined that exon 10 is skipped, producing a truncated protein (p.Gly412Alafs*86) (Figure S3). Furthermore, HEK.*SZT2*^c.1496G>T/KO^ cells displayed constitutive mTORC1 activity (Figure 1C and Figure S4). We generated homozygous *SZT2* p.Val1984del HEK cells (HEK.*SZT2*^p.Val1984del / p.Val1984del^) and determined that this VUS also resulted in constitutively active mTORC1 (Figure 1B-C). Collectively, these results demonstrate that complete *SZT2* LoF and both the previously published c.1496G>T and the recurrent p.Val1984del lead to constitutive mTORC1 signaling also suggesting *SZT2* LoF.

### A medium-throughput assay for functional characterization of *SZT2* VUS

The time-consuming process of limited dilution cloning and identification of clones with desired genotypes led us to investigate methods to increase throughput of our approach. Based on high rates of indel formation (60+%) due to nonhomologous end-joining (NHEJ) as well as high HDR efficiency (up to 30%) in HEK293T, we considered the possibility of performing functional testing directly after gene editing. Though immunoblot analysis from a pool of cells with different *SZT2* alleles would be challenging to interpret, we hypothesized that immunolabeling for phospho-S6 (P-S6) followed by flow cytometry would be a viable alternative. Amino acid starved *SZT2* null cells were shown to have both elevated P-S6K and P-S6 levels due to constitutive mTORC1 activity ^18^. The conceptual framework is as follows: if a variant is LoF variant, then cells homozygous for that variant or compound heterozygous for the variant and a truncating variant induced by NHEJ would lack any functional SZT2 and therefore exhibit constitutive mTORC1 activation. Alternatively, if a variant did not cause SZT2 LoF, cells with one or two copies would express sufficient levels of functional SZT2 for physiological regulation of mTORC1 by amino acid deprivation (Figure 2A). Visually, a single right-shifted peak (P-S6^HIGH^) would be indicative of a LoF variant, while observation of two peaks (i.e. P-S6^HIGH^ and P-S6^LOW^) would indicate the variant did not cause LoF. Subsequent genotyping of unsorted cells as well as the sorted pools of high and low phosphorylated-S6 (i.e. P-S6^HIGH^ and P-S6^LOW^) allowed us to determine the CMAS for each allele.

**Figure 2.**
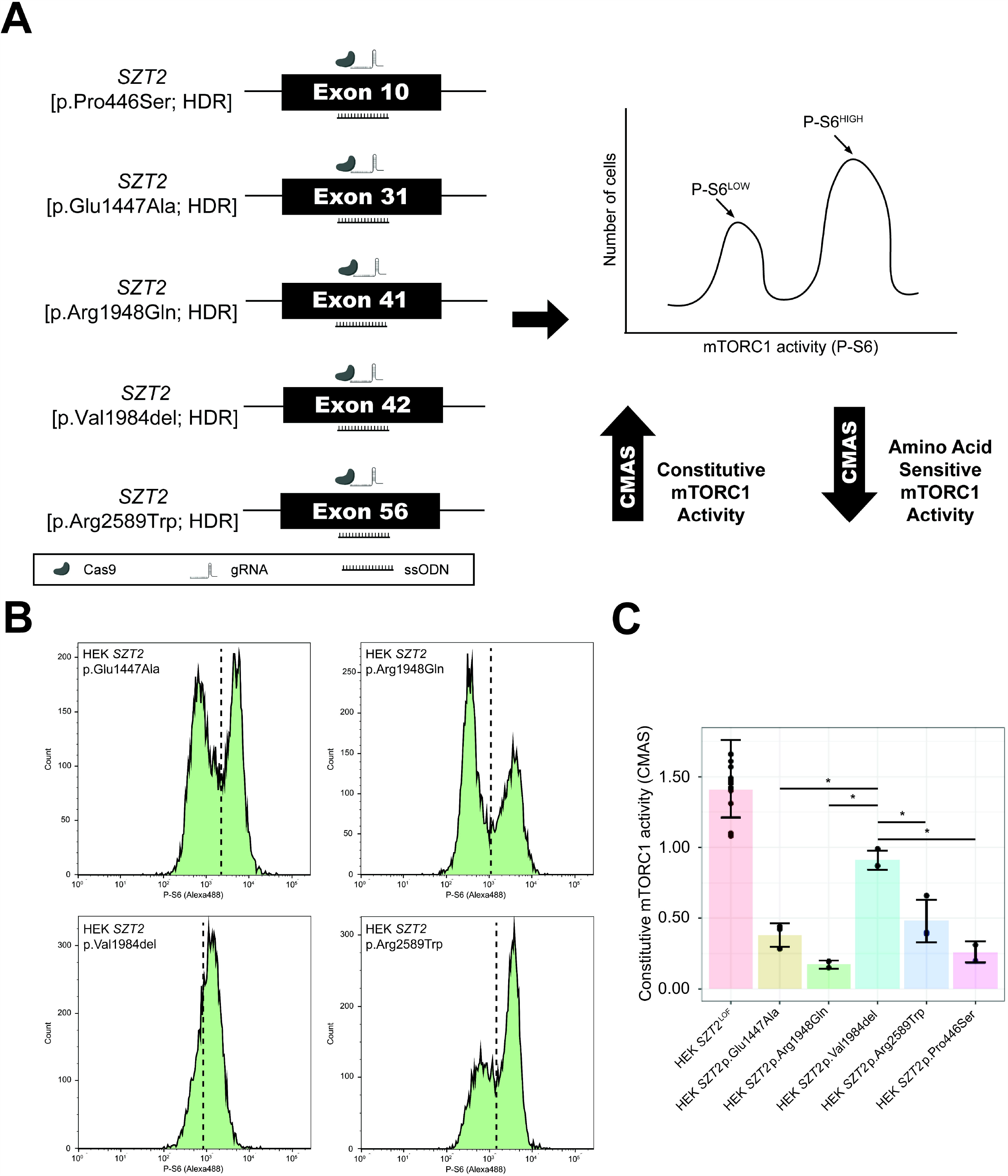
Development of medium throughput assay for functional characterization of *SZT2* VUSs. (A) HEK cells were co-transfected with px459 encoding targeting gRNAs and repair oligonucleotides, followed by puromycin selection. Cells were starved of amino acids and then fixed for immunolabeling of phosphorylated S6 and FACS sorting. gDNA was isolated from unsorted and sorted cells followed by amplicon sequencing to confirm CRISPR/Cas9 targeting and to calculate CMAS. (B) Representative FACS plots. Dashed line represents the boundary for sorting P-S6^HIGH^ and P-S6^LOW^ cell populations. (C) CMAS scores derived from amplicon sequencing of sorted and unsorted cells. For all but p.Arg1948Gln (n=2) and p.Pro466Ser (n=2), at least 3 individual replicates were performed. * = p < 0.05 (One-way ANOVA with Tukey’s posthoc test).

As a proof-of-principle, we first performed phospho-S6 FACS sorting on amino acid starved wildtype HEK and HEK *SZT2*^KO/KO^ cells. Wildtype HEK cells exhibited a single left-shifted peak (P-S6^LOW^), while the *SZT2*^KO/KO^ cells exhibited a single right-shifted peak (P-S6^HIGH^), indicating constitutive mTORC1 activity (Figure S5). We then used the assay to investigate a number of the *SZT2* VUSs present in the cohort: p.Val1984del, p.Glu1447Ala, p.Arg1948Gln, and p.Arg2589Trp (Figure 2B). As an additional negative control, we assayed a common variant in the general population, *SZT2* p.Pro446Ser (MAF=0.3 in gnomAD), and present in a homozygous state in multiple individuals (n=15,475). All LoF alleles had high CMAS scores (mean CMAS=1.41 ± 0.2; n=13) consistent with constitutively active mTORC1, while CMAS for *SZT2* p.Pro446Ser was low (0.26 ± 0.07; n=2), consistent with the protein retaining physiological scaffolding function and amino acid sensitive mTORC1 (Figure 2C and S6). As with the western blot analysis above, p.Val1984del was significantly enriched in the P-S6^HIGH^ (mean CMAS = 0.91 ± 0.06; n=3) pool, indicative of constitutive mTORC1 activity. Conversely, p.Glu1447Ala (mean CMAS = 0.38 ± 0.08), p.Arg1948Gln (mean CMAS = 0.17 ± 0.03; n=2), and p.Arg2589Trp (mean CMAS = 0.48 ± 0.15) were not enriched in the P-S6^HIGH^ pool, (Figure 2 B-C) and CMAS were not significantly different to p.Pro446Ser, but rather were consistent with the CMAS of the benign variant, suggesting these variants do not cause SZT2 LoF.

### Haplotype analysis of recurrent SZT2 in-frame deletion p.Val1984del

Based on our observation of p.Val1984del in multiple individuals, we hypothesized this variant may be derived from a common ancestor. In support of this observation, *SZT2* p.Val1984del is observed at low allele frequency in two populations in gnomAD, specifically in non-Finnish Europeans (MAF=0.00005420) and Ashkenazi Jewish (MAF=0.0008679), though no homozygous individuals are reported. Using exome sequencing data we identified a shared haplotype spanning ∼4 Mb including the *SZT2* p.Val1984del in all three individuals in our cohort (Figure 3), suggesting this variant is a founder variant in those of Jewish ancestry.

**Figure 3.**
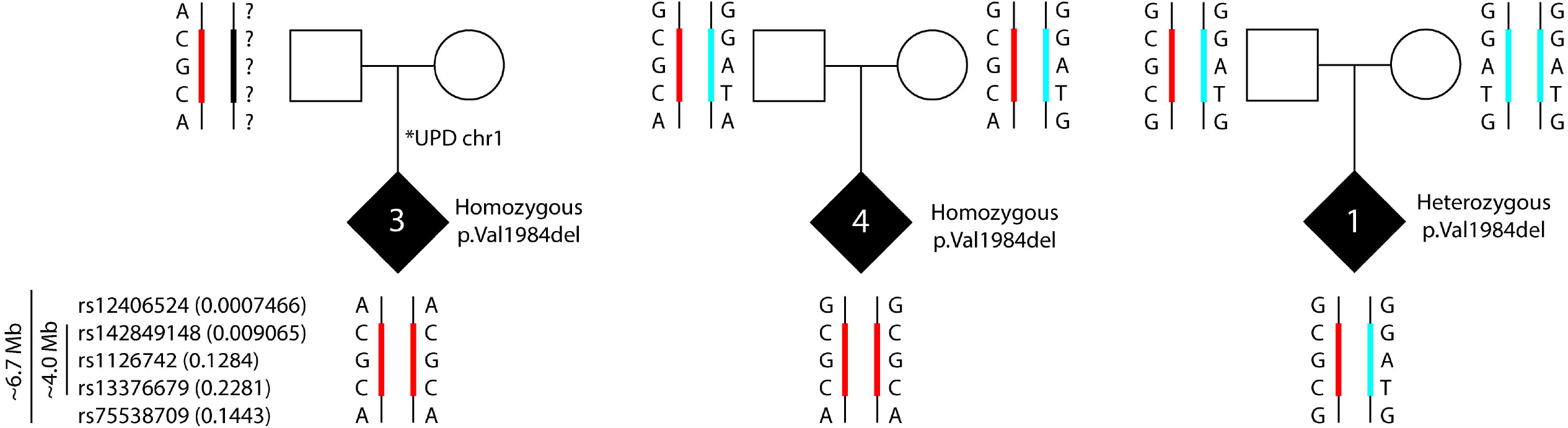
Shared haplotype suggests SZT2 p.Val1984del is a founder variant in those of Ashkenazi Jewish ancestry. Using exome sequencing data from three study trios in combination with allelic frequencies in the gnomAD population database, all individuals carry the same rare (MAF ranging from 0.00007-0.2) variants spanning a 4Mb interval. Ref|Alt alleles for SNVs (1 & 5 flanking; 2-4 in haplotype block): (1) rs12406524=G|A; (2) rs142849148=G|C; (3) rs1126742=A|G; (4) rs13376679=T|C; (5) rs75538709=G|A.

### Clinical characterization of individuals with bi-allelic SZT2 variants

The functional assays allowed us to tentatively reclassify 8/16 (50%) VUS in our cohort of 12 individuals. We acknowledge that reclassification of variants based on functional data is relevant in a clinical setting when classification is based on a ‘well-established assay’, which our approach has not yet achieved ^20^. However, for simplicity sake in this research study we reclassified VUS with CMAS similar to p.Pro446Ser (GnomAD MAF=0.3, numerous homozygotes) as likely benign (LB) and VUS with CMAS consistent with SZT2 LoF as likely pathogenic (LP). We then grouped these individuals into four groups (P or LP/P or LP, P or LP/VUS, VUS/VUS, LB/LB) and assessed the prevalence of the most common clinical features associated with pathogenic *SZT2* variants i.e. early-onset epilepsy, developmental delays, intractable focal seizures, macrocephaly and corpus callosum abnormalities (Table 1 and 2) ^2-16^. Individual 2 with bi-allelic *SZT2* truncating variants did not require reclassification but was included in the cohort as a representative of the P/P group for the purpose of phenotypic comparison. While sub-grouping the cohort in this manner meant we were unable to perform statistical analysis due to small group sizes, we could make a number of observations. The median seizure onset was 24 months and relatively consistent, with the exception of the LB/LB group where seizure onset was much later (median 13 years) and a number of the core features of *SZT2*-associated epilepsy were present in only one of the two individuals with these variants. Conversely, most individuals with either PLP/PLP or PLP/VUS did present with focal seizures, developmental delay and macrocephaly. Notably, where this information was available, the majority of individuals achieved seizure control, most with anti-seizure medications, but some were self-limiting (Table S4). Moreover, very few individuals, except individuals 9 and 11, had corpus callosum abnormalities (Table S5 and Figure S6).

**Table 2.** Summary of clinical features stratified by ACMG criteria and functional classification.

| Category | P or LP/<br>P or LP | P or LP/VUS | VUS/VUS | LB/LB | Cohort |
| --- | --- | --- | --- | --- | --- |
| Affected individuals | 1-4 (n=4) | 5-8 (n=4) | 9,10 (n=2) | 11,12 (n=2) | **n=12** |
| Median seizure onset (range) | 30m (2m-4y) | 9m (2DOL-2y) | 18m (3DOL-3 y) | 13 y (6y10m-20y) | **24m (2DOL-20y)** |
| Focal seizure | 3/4 (75%) | 2/4<sup>&</sup> (50%) | 0/2 (0%) | 1/2 (50%) | **6/12 (50%)** |
| Seizures intractable | 2/4 (50%) | 1/3 (33%) | 0/2 (0%) | 1/2* (50%) | **4/11 (36%)** |
| Developmental delays | 4/4 (100%) | 4/4 (100%) | 2/2 (100%) | 1/2 (50%) | **11/12 (92%)** |
| Macrocephaly | 3/4<sup>^</sup> (75%) | 2/4<sup>%</sup> (50%) | 1/2 (50%) | 1/2 (50%) | **7/12 (58%)** |
| Corpus callosum abnormalities | 0/4 (0%) | 1/4 (25%) | 1/2 (50%) | 1/2 (50%) | **3/12 (25%)** |

<sup>^</sup> Individual 4 had microcephaly <sup>&</sup> individual 7 had no report of seizures and <sup>%</sup> microcephaly <sup>\*</sup>individual 12 achieved only partial seizure control Abbreviations: DOL, day of life; m, months; y, years

## Discussion

The identification of VUS is one of the most challenging bottlenecks in clinical genetic diagnostics of the modern era. We developed a novel individualized platform that allowed us to recategorize 8/16 (50%) of *SZT2* VUS. Of note, we identify a recurrent in-frame deletion (p.Val1984del) in four individuals, including two in a homozygous state (one instance of chromosome 1 UPD), and one heterozygous in trans with a truncating variant. Two of the individuals (1 and 3) were identified at Lurie Children’s hospital over two years. We determined that this variant is a founder variant in individuals with Jewish ancestry. A carrier frequency of 1:576 is estimated based on gnomAD data, but may differ on local ancestry, and inclusion of this variant in genetic testing panels for those with Jewish ancestry requires further investigation.

Based on the necessity for a functional assay to reclassify *SZT2* VUSs, we developed a strategy to functionally characterize *SZT2* VUSs by knock-in of *SZT2* variants into HEK 293T cells. The molecular function of *SZT2* as a critical regulatory scaffolding protein in the amino acid sensing arm of the mTOR signaling pathway had previously been elucidated in HEK 293T ^17; 18^. Advantages of using HEK 293T cells are the high transfection and gene editing efficiencies, both for knockout (NHEJ) and knock-in (HDR) strategies. We successfully developed this strategy as a medium-throughput assay that revealed a recurrent inframe deletion *SZT2* p.Val1984del to be a LoF variant (Figure 2B-C and Figure 3B-C). Using this FACS-based assay we also functionally characterized an additional set of 3 SZT2 VUSs (p.Glu1447Ala, p.Arg1948Gln, and p.Arg2589Trp) as unlikely to be LoF as these variants retained amino-acid sensitive mTORC1 activity as observed for a common population variant (p.Pro446Ser) and suggest these variants are likely benign. However, while our approach can robustly detect complete LoF alleles, it may not detect more subtle effects on protein function, including partial LoF. Moreover, we know that the clinical features of individuals with pathogenic *SZT2* variants are neuronally restricted, even though *SZT2* is ubiquitously expressed, thus perturbation of the mTOR pathway is more likely to have a detrimental impact during neuronal development and/or function. For instance, it has recently been shown that mTOR regulation is essential for outer radial glia migration during human neuronal development ^23^. In the future, development of high-throughput functional assays in cells of a neuronal lineage, for instance, immortalized ReNcells, may improve on the discriminatory power of the novel assay we present here.

Nakamura Y et al. recently examined mTORC1 activity in lymphoblastoid cell lines (LCLs) from affected individuals^24^. They report elevated mTORC1 activity in cell lines derived from individuals with biallelic *SZT2* variants relative to cell lines generated from healthy individuals. There are a few important caveats to their study. First, the developed assay requires generating LCLs from individuals, which is not always possible and decreases throughput. Further, both cell lines generated from individuals with biallelic *SZT2* LoF variants show significant response to amino acid treatment after starvation. One of the key findings from the initial studies, and our studies here in HEK293T cells, was that *SZT2* gene knockout rendered cells completely insensitive to amino acids ^17; 18^. Although difficult to explain, the finding could be a technical issue with the LCL model.

Similar to previously published biallelic *SZT2* cases, most individuals carrying either P/LP or VUS presented with pediatric-onset epilepsy and expressed common features including focal seizures, developmental delay and macrocephaly. However, none of the individuals in these groups had corpus callosum abnormalities, including those with P/LP variants, suggesting that this is not a cardinal feature of *SZT2*-associated epilepsy. Moreover, a genotype-phenotype correlation has been suggested; individuals bearing truncating mutations may be more likely to have intractable seizures than individuals with missense variants ^12^. In this cohort, we observed more variability, with individual 2 (bi-allelic truncations) having seizure onset at two months with intractable seizures, while the other individual with intractable seizures (with seizure onset at two days of life) carried a multi-exon in-frame deletion and a VUS (Table S4, S5). Moreover, the individuals with homozygous p.Val1984del (individuals 3, 4) had divergent presentations, with individual 3 presenting with DEE and intractable seizures while individual 4 had a suspected neonatal seizures that resolved without medication. Despite these differing seizure patterns, both individuals have developmental delays and cognitive impairment. Finally, one individual had adult-onset epilepsy with seizures eventually being well-controlled on anti-convulsant medication. This individual had dysgenesis of the corpus callosum (Figure S6) and was found to have inherited *SZT2* missense VUS from each parent. This case is rather intriguing, as it potentially suggests the phenotypic spectrum for biallelic *SZT2* variants is significantly broader than previously thought. However, testing of these variants utilizing our assay suggests they do not cause SZT2 LoF, although we cannot rule out the possibility of partial LoF variants, as described above, which could account for the milder phenotypic presentation. Collectively, these results suggest a straightforward genotype-phenotype relationship is unlikely and further studies are needed to characterize the phenotypic spectrum and the impact of genetic variation on protein function.

In summary, here we demonstrate the utility of an individualized platform to recharacterize *SZT2* VUS. Importantly, this included a p.Val1984del variant that has a carrier allele frequency of at least 1:576 in those of Jewish ancestry and is a founder variant in this population. While additional modifications are still required to increase throughput, perhaps using saturation mutagenesis or multiplex assays of variant effect (MAVE) ^25; 26^, our approach can be applied to characterize VUS in other mTORopathies, including T*SC1, TSC2, DEPDC5, NPRL2* and *NPRL3*, which are the most common causes of focal epilepsies. As the mTORopathies are the targets of multiple new clinical trials for mTORC inhibitors, resolution of VUSs could qualify more individuals with intractable epilepsies for inclusion in these studies ^27^.

## Supplemental Data

Supplemental data includes seven figures and five tables.

## Acknowledgments

This work was sponsored by NIH NINDS R00NS089858 (GLC). We thank the NUCATS TL1 award for support (TR001423 to JDC). We thank Dr. Suchitra Swaminathan and Paul Mehl for assistance with flow cytometry. This work was supported by the Northwestern University – Flow Cytometry Core Facility supported by Cancer Center Support Grant (NCI CA060553). Flow Cytometry Cell Sorting was performed on a BD FACSAria SORP system and BD FACSymphony S6 SORP system, purchased through the support of NIH 1S10OD011996-01 and 1S10OD026814-01.

## Web Resources

gnomAD v2.1.1 (https://gnomad.broadinstitute.org) - last accessed 23 Oct 2020

SeattleSeq (https://snp.gs.washington.edu/SeattleSeqAnnotation138/) – last accessed 22 Oct 2020

Clinvar (https://www.ncbi.nlm.nih.gov/clinvar/) - last accessed 22 Oct 2020

