## Supplemental Material for "Determining the pathogenicity of variants of uncertain significance and identification of a founder variant in the epilepsy-associated gene, *SZT2*"

**SUPPLEMENTAL DATA**

**Table of contents**

| Supplemental materials and methods | Page 3 |
| --- | --- |
| Figure S1: Distribution of variants throughout *SZT2*. | Page 3 |
| Figure S2: Amino acid treatment of HEK *SZT2*^KO/KO^ cells. | Page 3 |
| Figure S3: Exon skipping in *SZT2* transcript caused by c.1496G>T. | Page 4 |
| Figure S4: Amino acid treatment of HEK *SZT2*^c.1496G>T/KO^ cells. | Page 4 |
| Figure S5: Flow cytometry based assay for mTORC1 activity in amino acid starved cells. | Page 5 |
| Figure S6: Flow cytometry based assay for mTORC1 activity in amino acid starved HEK *SZT2* p.Pro446Ser cells. | Page 6 |
| Figure S7: MRI images in individuals 10 and 11. | Page 6 |
| Table S1: *SZT2* gRNAs and primers. | Page 7 |
| Table S2: Antibodies. | Page 8 |
| Table S3: Percentage of alleles determined by amplicon sequencing in unsorted and P-S6 sorted cells. | Page 9 |
| Table S4: Seizure characteristics and epilepsy diagnosis in individuals with biallelic *SZT2* variants | Page 10 |
| Table S5: Developmental History and Other Features in individuals with biallelic *SZT2* variants | Page 11 |
| References | Page 13 |

**Supplemental materials and methods**

*Exon skipping PCR*

RNA was extracted from HEK 293T cells with Trizol (Invitrogen 15596026) according to manufacturer’s recommendations. cDNA was synthesized using iScript (Biorad) and 1000 ng of input RNA according to manufacturer’s recommendations. Primers amplifying from *SZT2* exon 9 to exon 11 were used to generate amplicons (5’ aatgagcacctggtctctgc 3’ and 5’ ggcactgaggagaaggactg 3’). Amplicons were separated on 2% agarose followed by gel purification and Sanger sequencing (Figure S3).

**Supplementary Figures**

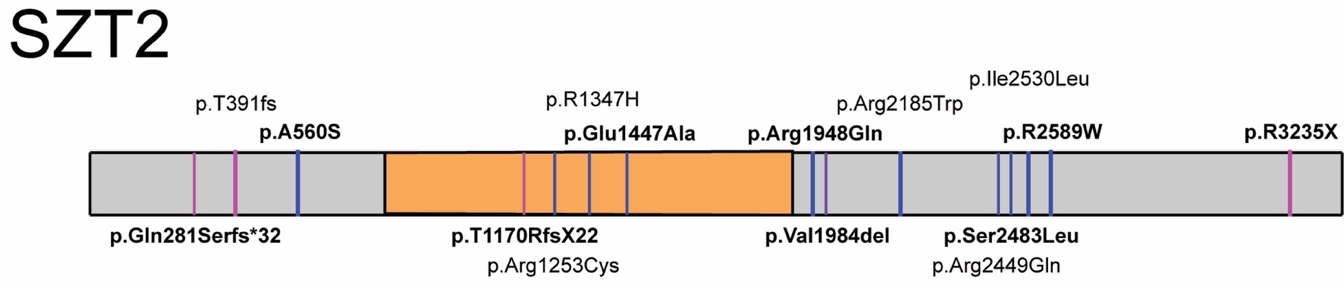

**Figure S1: Distribution of variants throughout *SZT2***. Location of truncation variants indicated by purple line, while missense or in-frame deletions are represented by blue lines. The multi-exon deletion is displayed in orange.

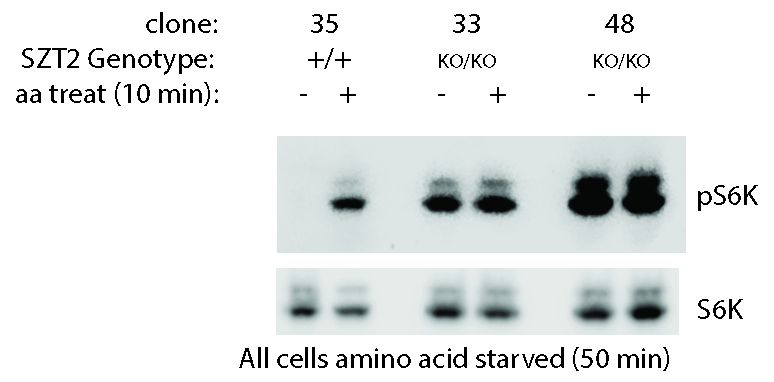

**Figure S2: Amino acid treatment of HEK *SZT2*^KO/KO^ cells.** *SZT2*^KO/KO^ cells were generated using a gRNA (GTGGCAGCCAGATGAACCAG) targeting exon 3 as previously described followed by puromycin treatment and limited dilution cloning to establish clonal lines (1). We observed excess mTORC1 activity in amino acid starved *SZT2*^KO/KO^ cells relative to control *SZT2*^+/+^ cells. AA: - denotes cells starved of amino acids for 60 min; AA: + denotes cells starved of amino acids for 50 min followed by subsequent treatment with amino acids for 10 min. pS6K = phosphorylated S6K. For genotype, + = wildtype or reference allele.

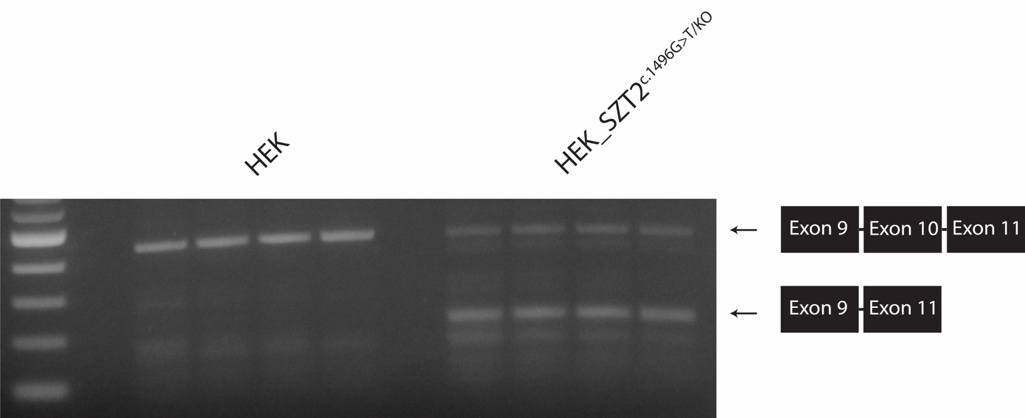

**Figure S3: Exon skipping in *SZT2* transcript caused by c.1496G>T.** Sanger sequencing of smaller PCR products in cells heterozygous for *SZT2* c.1496G>T confirmed exon skipping due to disruption of splice donor site. The four lanes of control HEK (HEK_SZT2^WT^/^WT^) and four lanes of compound heterozygous HEK_SZT2^c.1496G>T/KO^ each contain two biological replicates and two technical replicates.

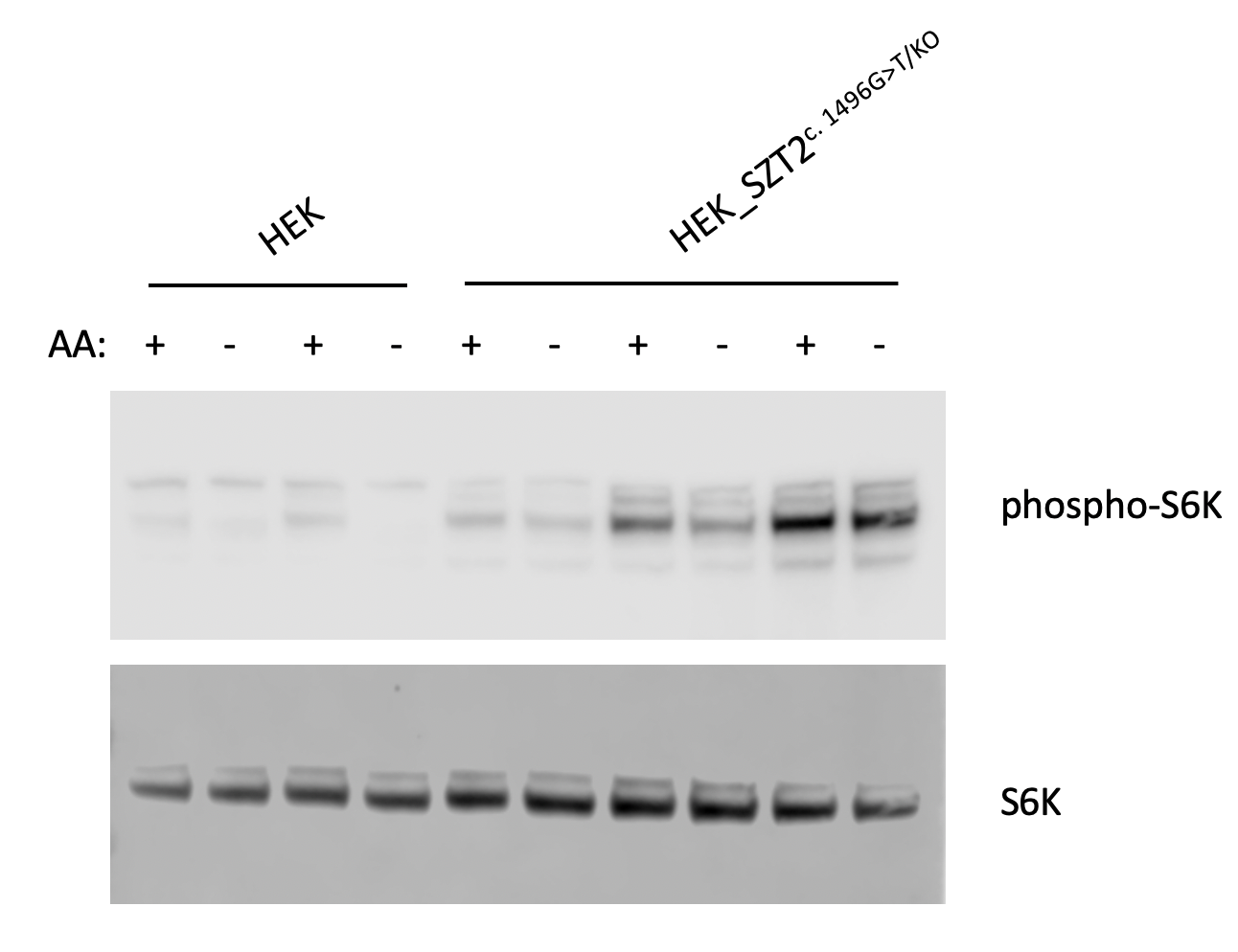

**Figure S4: Amino acid treatment of HEK *SZT2*^c.1496G>T/KO^ cells.** *SZT2*^c.1496G>T/KO^ cells exhibit excessive mTORC1 activity under amino acid starvation, in support of *SZT2* c.1496G>T as a loss-of-function allele. Two biological replicates shown for control HEK (HEK_SZT2^WT^/^WT^). Three biological replicates are shown for compound heterozygous HEK_SZT2^c.1496G>T/KO^. AA: - denotes cells starved of amino acids for 60 min; AA: + denotes cells starved of amino acids for 50 min followed by subsequent treatment with amino acids for 10 min.

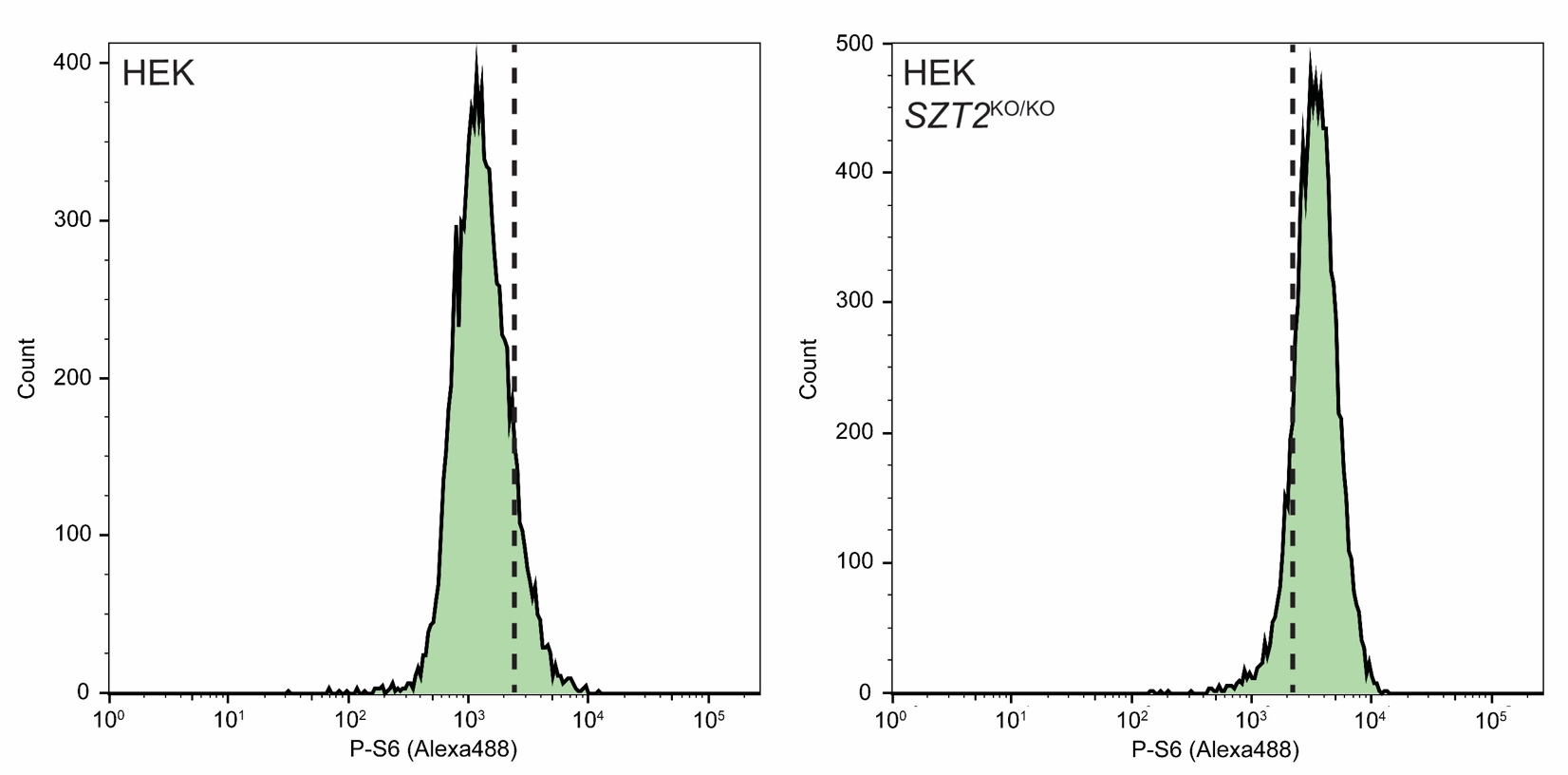

**Figure S5: Flow cytometry based assay for mTORC1 activity in amino acid starved cells.** P-S6 levels are higher in amino acid starved *SZT2*^KO/KO^ cells relative to control HEK cells due to constitutive mTORC1 activity. Dashed line denotes the gating cutoff used to separate P-S6^LOW^ from P-S6^HIGH^ cell populations.

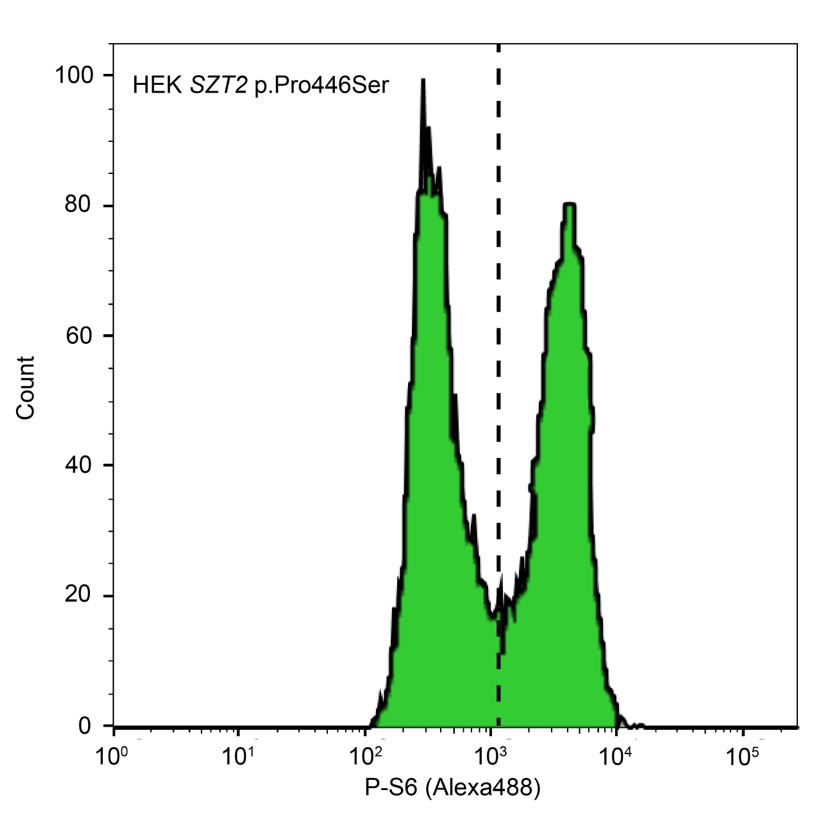

**Figure S6: Flow cytometry based assay for mTORC1 activity in amino acid starved HEK *SZT2* p.Pro446Ser cells.** We observed two clear cell populations, suggesting the SZT2 p.Pro446Ser variant does not lead to constitutive mTORC1 activation.

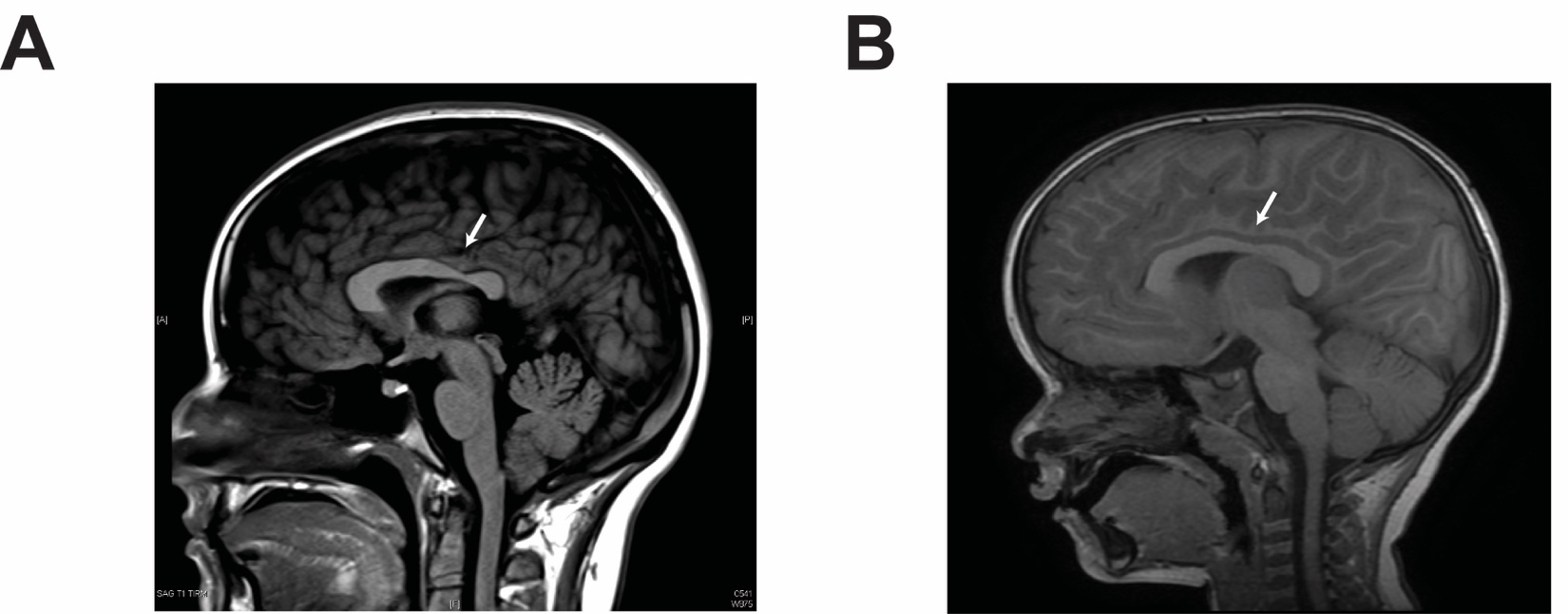

**Figure S7: MRI images from individuals 10 and 11.** Arrows indicating dysgenesis of corpus callosum in individual 11 (A) and possible corpus callosum abnormality (upper limit of normal) in individual 10 (B).

**Table S1: *SZT2* gRNAs and primers.**

| **Primer seq (5’ to 3’)** | **Name** | **Purpose** | **HDR/patient allele** |
| --- | --- | --- | --- |
| CACCGGTGGCAGCCAGATGAACCAG | SZT2_ex3s | gRNA pair | SZT2^KO/KO^  (exon 3) |
| AAACCTGGTTCATCTGGCTGCCACC | SZT2_ex3as |  |  |
| CACCGTTTCTGGAACACGCTGCAG | SZT2_cG1496Ts | gRNA pair | *SZT2* c.1496G>T |
| AAACCTGCAGCGTGTTCCAGAAAC | SZT2_cG1496Tas |  |  |
| CTCCCCAGCTCCCGCCTTTGCAAATCTGTCCCTCCATACAACAAATGGGGCAGCACTCATTGAGTGATGACTTCACTGACATCTGCAGCGTGTTCCAGAAACGCCGGATAACATGGGTACGATACAATGAACGAATGGGCTGCCTTAGTGCACAGGACACA | SZT2_cG1496TssODN | Repair oligo |  |
| TTGGAGGTAAAGCTGGTGCT | SZT2_cG1496TsurvF | Primer pair for T7eI assay / Sanger sequencing |  |
| GTCAGGAAGCGTGAAATGCT | SZT2_cG1496TsurvR |  |  |
| TCGTCGGCAGCGTCAGATGTGTATAAGAGACAGGGGAGGTAAGGGTGGTGAGT | SZT2_cG1496TampSeqF | Primer pair for amplicon sequencing |  |
| GTCTCGTGGGCTCGGAGATGTGTATAAGAGACAGCCTGTGGGTGTGTCCTCTGT | SZT2_cG1496TampSeqR |  |  |
| CACCGTTCTCCTGTGACGTTGTGTG | SZT2_1984DELs | gRNA pair | *SZT2* p.Val1984del |
| AAACCACACAACGTCACAGGAGAAC | SZT2_1984DELas |  |  |
| ATGCCAATCACATACCCCGAGAGACCCCCATGCTGGGCCCCATTTTGAGGCGTGAATGGACTCGGATCACAGTTCCCCACACGTCACAGGAGAAATGGCCAGGGACAAACTGCATTGTGGCTGCCAGGTACCCACGGGGCGCACAGCTCTCATCAGCAGC | SZT2_1984DELssODN | Repair oligo |  |
| TTGCCTGCCTTGATACCTCT | SZT2_1984DELsurvF | Primer pair for T7eI assay / Sanger sequencing |  |
| TTTCCAGATCCTTCCAATGC | SZT2_1984DELsurvR |  |  |
| TCGTCGGCAGCGTCAGATGTGTATAAGAGACAGAGGGTGGTGTGTCCCATTT | SZT2_1984DELampSeqF | Primer pair for amplicon sequencing |  |
| GTCTCGTGGGCTCGGAGATGTGTATAAGAGACAGATCCTTCCAATGCTCACCTG | SZT2_1984DELampSeqR |  |  |
| CACCGAGACACATCTGCCTGCTGTG | SZT2_1447s | gRNA pair | *SZT2* p.Glu1447Ala |
| AAACCACAGCAGGCAGATGTGTCTC | SZT2_1447as |  |  |
| GCGAGGCAGAGTCTAGCCCAGCAGGCCCCAGGTCTGATTCACGGCTCTCCCGGTATTCTACCTCTAGCTCTGGGTCACTCGCAGTGACGACACAGCAGGCAGATGTGTCTCCTAGTAGTGCATGTCACATGGTCAATGAGAGAGGCTGGCAGTGGAACCCT | SZT2_1447ssODN | Repair oligo |  |
| CCTGTTGAGGGGTGACTGAC | SZT2_1447survF | Primer pair for T7eI assay / Sanger sequencing |  |
| CACTTAGGCAGGCTGTTCCA | SZT2_1447survR |  |  |
| TCGTCGGCAGCGTCAGATGTGTATAAGAGACAGGCCGAGATTCAAGGGTTCCA | SZT2_1447ampSeqF | Primer pair for amplicon sequencing |  |
| GTCTCGTGGGCTCGGAGATGTGTATAAGAGACAGGTCTACGTCTGACAGCGAGG | SZT2_1447ampSeqR |  |  |
| CACCGTCCCTTCCACTCCCGTCAGC | SZT2_1948s | gRNA pair | *SZT2* p.Arg1948Gln |
| AAACGCTGACGGGAGTGGAAGGGAC | SZT2_1948as |  |  |
| TCGGTAAAAGAAAGGGCGGCAGCACTGCCTTGTGAGGGAGGCTCCTGGGTGGGATCTCACCATCACTGGGCAGTGGTGCCTGCTGACGGCTGTGGAAGGGAGTCTCACTGCGCCACAGATCTTCTTCACTCTCGGCCACCAGAAGAGAGTTACACACGTGG | SZT2_1948ssODN | Repair oligo |  |
| GACCACCACCACCCACTTAG | SZT2_1948survF | Primer pair for T7eI assay / Sanger sequencing |  |
| CCTTGCCCCCATCTTCCTTT | SZT2_1948survR |  |  |
| TCGTCGGCAGCGTCAGATGTGTATAAGAGACAGAGACCTTCATGACAGCCACG | SZT2_1948ampSeqF | Primer pair for amplicon sequencing |  |
| GTCTCGTGGGCTCGGAGATGTGTATAAGAGACAGACTCTGCCCTCAGCTTTCAC | SZT2_1948ampSeqR |  |  |
| CACCGGCGAAATGCTCCCCGGCAG | SZT2_2589s | gRNA pair | *SZT2* p.Arg2589Trp |
| AAACCTGCCGGGGAGCATTTCGCC | SZT2_2589as |  |  |
| ACGGAACCCCTCTTCCCACCCAGTCCCTGGCCCCATATTTACCTTCTTGTCCACAACCTCTAGTAGCAAGAGTCTCTGCCAGGGAGCATTTCGCCCTGAGCTCCCATCACCTCCTGGCTCGAAGCGCTGCATGGCTTTGGCAGCTACAGGGTGGGGGAGGG | SZT2_2589ssODN | Repair oligo |  |
| AGATCTTCGGCCCTTGTTCC | SZT2_2589survF | Primer pair for T7eI assay / Sanger sequencing |  |
| CCTTAGGGCCTCGTCAAAGG | SZT2_2589survR |  |  |
| TCGTCGGCAGCGTCAGATGTGTATAAGAGACAGATCTGTTCCCAGGCCCCTAT | SZT2_2589ampSeqF | Primer pair for amplicon sequencing |  |
| GTCTCGTGGGCTCGGAGATGTGTATAAGAGACAGTGAGATCACGGAACCCCTCT | SZT2_2589ampSeqR |  |  |
| CACCGAGTATGTGGCTATGGCACCC | SZT2_446s | gRNA pair | *SZT2* p.Pro446Ser |
| AAACGGGTGCCATAGCCACATACTC | SZT2_446as |  |  |
| ACATCATGCAAAATGTCGTAGCCGCCTTCCATCGTCACTTCCACCCGTGTTACTCGAGGGCCCTCAGGCTCCAGGGGCCAGGAAGCCATAGCCACATACTCAATGCGCATGTTGTGTTTCCACAGCAGCACCAGCTTTACCTCCAATTGGGACCCTCCTT | SZT2_446ssODN | Repair oligo |  |
| TTGGAGGTAAAGCTGGTGCT | SZT2_446survF | Primer pair for T7eI assay / Sanger sequencing |  |
| GTCAGGAAGCGTGAAATGCT | SZT2_446survR |  |  |
| AGGTAGGTGGGGGTTTCAGA | SZT2_446ampSeqF | Primer pair for amplicon sequencing |  |
| ATGCAAAATGTCGTAGCCGC | SZT2_446ampSeqR |  |  |

Abbreviations: s = sense, as = antisense, ssODN = single strand oligo donor, surv = surveyor, ampSeq = amplicon sequencing, F = forward, R = reverse. Amplicon sequencing primers were synthesized with standard Illumina adaptors.

**Table S2: Antibodies.**

| **Primary antibodies** | | **Secondary antibodies** | |
| --- | --- | --- | --- |
| **Vendor information** | **Binding condition** | **Vendor information** | **Binding condition** |
| Cell Signaling Technologies (CST) rabbit anti-phospho-S6K (108D2) | 1:1000 overnight at 4 deg C (Western blot) | Abcam HRP-conjugated goat anti-rabbit (ab205718) | 1:50,000 room temperature for 1 hr |
| CST rabbit anti-S6K (49D7) | 1:1000 overnight at 4 deg C (Western blot) | Abcam HRP-conjugated goat anti-rabbit (ab205718) | 1:50,000 room temperature for 1 hr |
| CST Alexa488 conjugated rabbit anti-phospho-S6 (#5018; D68F8) | 1:50 30 min at 4 deg C (labeling for FACS) | N/A | N/A |

**Table S3: Percentage of alleles determined by amplicon sequencing in unsorted and P-S6 sorted cells.**

| **Alleles (Replicate 1):** | **HDR (1)** | | | **LoF (1)** | | | **Ref (1)** | | |
| --- | --- | --- | --- | --- | --- | --- | --- | --- | --- |
| **Cell pool:** | **Unsorted** | **P-S6^LOW^** | **P-S6^HIGH^** | **Unsorted** | **P-S6^LOW^** | **P-S6^HIGH^** | **Unsorted** | **P-S6^LOW^** | **P-S6^HIGH^** |
| ***SZT2* p.Glu1447Ala** | 20.65 | 26.99 | 9.03 | 36.5 | 25.39 | 50.94 | 9.61 | 14.99 | 3.39 |
| ***SZT2* p.Arg1948Gln** | 20.64 | 28.91 | 3.14 | 5.98 | 4.22 | 9.93 | 4.76 | 5.99 | 0.37 |
| ***SZT2* p.Arg2589Trp** | 19.03 | 26.29 | 7.67 | 5.92 | 4.35 | 9.3 | 2.12 | 3.1 | 0.25 |
| ***SZT2* p.Val1984del** | 25.91 | 29.17 | 25.56 | 40.17 | 26.34 | 43.77 | 4.63 | 21.44 | 0.77 |
| ***SZT2* p.Pro446Ser** | 29.08 | 43.78 | 5.95 | 21.72 | 14.18 | 30.84 | 0.63 | 0.57 | 0 |
| **Alleles (Replicate 2):** | **HDR (2)** | | | **LoF (2)** | | | **Ref (2)** | | |
| **Cell pool:** | **Unsorted** | **P-S6^LOW^** | **P-S6^HIGH^** | **Unsorted** | **P-S6^LOW^** | **P-S6^HIGH^** | **Unsorted** | **P-S6^LOW^** | **P-S6^HIGH^** |
| ***SZT2* p.Glu1447Ala** | 22.5 | 26.27 | 9.44 | 31.26 | 24.87 | 50.37 | 8.73 | 11.19 | 2.99 |
| ***SZT2* p.Arg1948Gln** | 17.61 | 27.34 | 3.42 | 7.45 | 6.31 | 9.76 | 0 | 0.46 | 0 |
| ***SZT2* p.Arg2589Trp** | 14.75 | 24.69 | 9.76 | 5.76 | 3.78 | 8.57 | 0 | 0 | 0 |
| ***SZT2* p.Val1984del** | 23.76 | 34.29 | 20.67 | 39.15 | 28.93 | 42.88 | 0.4 | 1.72 | 0 |
| ***SZT2* p.Pro446Ser** | 29.51 | 43.68 | 9.18 | 20.59 | 14.46 | 30.17 | 0.31 | 0.49 | 0 |
| **Alleles (Replicate 3):** | **HDR (3)** | | | **LoF (3)** | | | **Ref (3)** | | |
| **Cell pool:** | **Unsorted** | **P-S6^LOW^** | **P-S6^HIGH^** | **Unsorted** | **P-S6^LOW^** | **P-S6^HIGH^** | **Unsorted** | **P-S6^LOW^** | **P-S6^HIGH^** |
| ***SZT2* p.Glu1447Ala** | 25.11 | 30.1 | 7.09 | 35.31 | 25.85 | 58.62 | 7.66 | 10.77 | 1.98 |
| ***SZT2* p.Arg1948Gln** |  |  |  |  |  |  |  |  |  |
| ***SZT2* p.Arg2589Trp** | 22.09 | 31.4 | 8.54 | 6.36 | 3.87 | 9.23 | 0 | 0 | 0 |
| ***SZT2* p.Val1984del** | 25.51 | 35.66 | 22.21 | 45.71 | 33.24 | 49.58 | 0.69 | 3.31 | 0 |
| ***SZT2* p.Pro446Ser** |  |  |  |  |  |  |  |  |  |

Abbreviations: Ref= Reference. For LoF variants, percentage of only the most abundant variant is reported.

**Table S4: Seizure characteristics and epilepsy diagnosis in individuals with biallelic *SZT2* variants**

| **Affected individual** | **1** | **2** | **3** | **4** | **5** | **6** | **7** | **8** | 9 | **10** | **11** | **12** |
| --- | --- | --- | --- | --- | --- | --- | --- | --- | --- | --- | --- | --- |
| Diagnosis | DEE | DEE | DEE | Suspected neonatal seizure | DEE | Infantile epilepsy | DEE | Infantile epilepsy | Suspected neonatal seizure | Infantile epilepsy | Focal epilepsy | DEE |
| Age at sz onset | 2 y | 2 m | 4 y | 3 y | 2 DOL | 2 y | No seizures* | 9 m | 3 DOL | 3 y | 20 y | 6 y |
| Current Age Sex | 5y, M | 8y, M | 6y, F | 10y; M | 7y, M | 5y9m; F | 5y10m; M | 10y; F | 6y; M | 5y, M | 23y, F | 9y; M |
| Febrile Seizures? | Yes | No | No | Unk | No | No | No | Yes | Unk | Yes | N | No |
| Seizure type(s) | Focal Impaired Awareness,  Focal to bilateral tonic-clonic | Focal Impaired Awareness,  Focal to bilateral tonic-clonic | Focal Impaired Awareness,  Focal to bilateral tonic | Generalized Tonic | Focal Impaired Awareness,  Focal to bilateral tonic | GTCS (rare) | NA | Atonic,  Absence,  Focal motor,  Generalized clonic | Generalized | Absence,  Generalized Tonic | Focal Impaired Awareness,  Focal to bilateral tonic-clonic | Myoclonic, Absence, Generalized |
| Current seizure control | Intractable | Intractable | Well controlled | Seizures resolved without medication | Intractable | Well controlled | N/A | Well controlled | Seizures resolved without medication | Well controlled | Partial | Intractable |
| Hx of status epilepticus? | Yes | Yes | Yes | N/A | Yes | No | N/A | No | N/A | No | No | No |
| Effective AED | CLB, OXC | CBD, ESM, LEV, ZNS, VNS (partially effective) | CLB, VPA | N/A | BRV, LCS, VPA | OXC | N/A | CLB, LEV, | N/A | LEV | LEV | CLB, VPA |
| Ineffective AEDs | VPA | ACZ, BRV, CLB, CLZ, FFM, LCS, LTG, OXC, PER, PB | LEV, | N/A | ACZ, CBD, ESM, FBM, FFM, LCS, LEV, OXC, TPM, PHT, CLB, VPA | N/A | N/A | CLZ, LTG,  TPM | N/A | OXC | N/A | N/A |
| Other ineffective therapies | N/A | KD | N/A | N/A | KD, MPN, VNS | N/A | N/A | N/A | N/A | N/A | Epidermoid cyst resection | N/A |

ACZ, acetazolamide; CBD, cannabadiol; CLB, clobazam; CLZ, clonazepam; DOL – day of life; ETX, ethosuximide; ESM, eslicarbamazepine; FBM, felbamate; FFM, rufinamide; GTCS, generalized tonic clonic seizure; KD, ketogenic diet; LEV, levetiracetam; LTG, lamotrigine; LCS, lacosamide; OXC, oxcabarbazepine; PER, perampanel; m, months, MPN, methylprednisolone; N/A, not applicable; PB, phenobarbital; PHT, phenytoin; Unk, uknown; VNS, Vagal Nerve Stimulator; VPA, valproate; TPM, topiramate; y, years; ZNS, zonisamide.

* This patient did not show evident motor seizures but his EEG was characterized by severe abnormalities consistent with encephalopathy (slow abnormalities and centro-parietal irritative elements)

**Table S5 Developmental History and Other Features in individuals with biallelic *SZT2* variants**

| **Affected individual** | **1** | **2** | **3** | **4** | **5** | **6** | **7** | **8** | 9 | **10** | **11** | **12** |
| --- | --- | --- | --- | --- | --- | --- | --- | --- | --- | --- | --- | --- |
| Developmental delay  /regression? | DD:  Walked at 21 m (requires AFOs), Non-verbal (uses sign-language) | DD:  Walked at 3 y (requires walker), Non-verbal | DD:  Walked at 15 m independently, Minimally verbal (uses communication device) | DD: Motor and speech regression reported during first year. Does not sit or talk, no eye contact | DD with regression at 7 y Walked at 2y but unable to walk after 7y, Non-verbal (uses sign language) | DD: Motor & speech regression at 2y4m. No crawling, no walking, only sitting & holding upright with assistance Non-verbal | DD:  Non-verbal, Unable to walk, feeding issues | DD:  Walked at 2y, in main-stream school with some assistance | DD: Not walking at 2.5 y, global DD with cognitive impairment | DD: Independent steps at 20m, fine motor delay, Non-verbal at 20m | No: Normal development | Developmental regression (speech& motor). Normal walking, Speech & cognitive challenges noted at 12m |
| Other neurological features | ASD | Hypotonia, ASD | ASD | Mixed muscle tone  Central hypotonia, hypertonia of mainly the left upper limb  No social contact | Hypotonia, hypertonia / spasticity, Unilateral conductive hearing loss, Self-injurious behavior | Stereotyped behavior | Hypotonia, Stereotyped movements | ADHD suspected | Hypotonia | Hypotonia, Astigmatism, and amblyopia, Poor social interaction | Epidermoid cyst (seizures persisted after resection), Depression | Intellectual disability (IQ 63) |
| Head circumference | Macrocephaly  (>98^th^ %ile) | Macrocephaly  (97.3 %ile) | Macrocephaly  (100%ile) | Microcephaly (-3 SD) | Macrocephaly  (>98^th^ %ile) | Macrocephaly,  (>99^th^ %ile) | Microcephaly (OFC 47 cm at 4 y, -2.11 SDS) | Normal | Macrocephaly  At 30 m 56 cm +4.5SD | Normal - 52cm at  3y 11m | Macrocephaly as child; Normal as adult (94  %ile) | Normal  52 cm at 7y10m  (41 %ile) |
| Dysmorphic features | None noted | Frontal bossing, Downslanted palpebral fissures, Hypertelorism (mild) | Wide nasal bridge | Down slanting palpebral fissures | Frontal bossing (mild), Flattened nasal bridge, Low set ears | Frontal bossing | Narrow forehead, Plagiocephaly, Hypotelorism, Single palmar crease | None noted | None noted | Frontal bossing, High arched palate | Large forehead,  Arched eyebrows | Protruding ears, Upslanting palpebral fissures, Thick eyebrows, Diastema |
| Dysgenesis of the corpus callosum | No | No | No | No | No | Yes - Slight narrowing of trunci corpori callosi | No | No | Yes - Thick corpus callosum | No* | Yes | No |
| Other  MRI findings | Non-specific diffusion restriction in the bilateral cerebellar hemispheres | Right PVNH & abnormal perisylvian gyral configuration;  Pineal cyst (13mm); Small pars intermedia cyst | None | Ischemic changes (s/p premature delivery and hemorrhage) | bilateral & multifocal areas of MCD involving both hemispheres with the appearance of PMG and/or associated deep sulci. particular frontal involvement, also suspected right parietal lobe involvement | No | Narrowing of the middle third of the Sylvian aqueduct (mild),  Myelination delay (mild) | Normal | None | None | Bilateral PVNH  Epidermoid cyst | None |

Abbreviations: AFO, Ankle foot orthotics; ADHD, attention deficit hyperactivity disorder; ASD, Autism spectrum disorder; DD, developmental delay; m, months; MCD, malformations of cortical development; MRI, Magnetic Resonance Imaging; PMG, polymicrogyria; PVNH, periventricular nodular heterotopia; y, years

* Noted as upper limit of normal’
